# Genetically-clustered antifungal phytocytokines and receptor proteins function together to trigger plant immune signaling

**DOI:** 10.1101/2023.11.27.568785

**Authors:** Julie Lintz, Yukihisa Goto, Kyle W. Bender, Raphaël Bchini, Guillaume Dubrulle, Euan Cawston, Cyril Zipfel, Sebastien Duplessis, Benjamin Petre

## Abstract

- Phytocytokines regulate plant immunity via cell-surface receptors. *Populus trichocarpa* RUST INDUCED SECRETED PEPTIDE 1 (PtRISP1) exhibits an elicitor activity in poplar, as well as a direct antimicrobial activity against rust fungi. *PtRISP1* gene directly clusters with a gene encoding a leucine-rich repeat receptor protein (LRR-RP), that we termed RISP- ASSOCIATED LRR-RP (PtRALR).
- In this study, we used phylogenomics to characterize the RISP and RALR gene families, and functional assays to characterize RISP/RALR pairs.
- Both *RISP* and *RALR* gene families specifically evolved in Salicaceae species (poplar and willow), and systematically cluster in the genomes. Two divergent RISPs, PtRISP1 and *Salix purpurea* RISP1 (SpRISP1), induced a reactive oxygen species (ROS) burst and mitogen-activated protein kinases (MAPKs) phosphorylation in *Nicotiana benthamiana* leaves expressing the respective clustered RALR. PtRISP1 triggers a rapid stomatal closure in poplar, and both PtRISP1 and SpRISP1 directly inhibit rust pathogen growth.
- Altogether, these results suggest that plants evolved phytocytokines with direct antimicrobial activities, and that the genes coding these phytocytokines co-evolved and physically cluster with their cognate receptors.

## Introduction

The plant immune system fends off pathogens and prevents diseases (Ngou *et al*., 2022). This system notably uses defense peptides commonly exhibiting either an antimicrobial activity or an immunomodulatory activity (Tavormina *et al*., 2015). Antimicrobial peptides (AMPs) possess a cytotoxic activity that targets and directly kills microbes (Bakare *et al*., 2022), whereas immunomodulatory peptides (also called phytocytokines analogous to metazoan cytokines) modulate cell immune signaling by binding to specific cell-surface receptors (Yamaguchi & Huffaker, 2011; Hou *et al*., 2021; Rhodes *et al*., 2021; Rzemieniewski & Stegmann, 2022). In animals, most defense peptides described to date are bi-functional, i.e., they exhibit both antimicrobial and immunomodulatory activities. These peptides are referred to as host defense peptides (HDPs), and emerge as molecules with high valorization potential, as their many activities can be exploited for therapeutic purposes (Yeung *et al*., 2011; Hilchie *et al*., 2013; Haney *et al*., 2019; Sun *et al*., 2023). In plants, only a handful of HDP candidates has been described and the concept of defense peptides having two distinct roles within the immune system has only recently emerged (Petre, 2020).

Plant cell-surface immune receptors reside at the plasma membrane and belong to either the receptor kinase (RK) or the receptor protein (RP) gene families (DeFalco & Zipfel, 2021). Members of those families with extracellular leucine-rich repeat (LRR) domains are referred to as LRR-RKs or LRR-RPs, and recognize peptide or protein ligands to initiate immune signaling events via the activation of intracellular kinases (Ngou *et al*., 2022). Unlike LRR-RKs, LRR-RPs lack a cytosolic kinase domain and require the universal adaptor kinase SUPPRESSOR OF BIR11 (SOBIR1) to accumulate and initiate immune signaling (Liebrand *et al*., 2013; Bi *et al*., 2016; Gust *et al*., 2017; Ranf, 2017). Immune receptor activation rapidly triggers a set of downstream responses; notably the transient accumulation of reactive oxygen species (ROS) and the activation by phosphorylation of mitogen-activated protein kinases (MAPKs) (Gust & Felix, 2014). Among the 19 LRR-RPs characterized as immune receptors so far, only the INCEPTIN RECEPTOR (INR) recognizes a plant peptide (Snoeck *et al*., 2023).

The Salicaceae family of plants regroups two main genera: *Populus* (poplar trees) and *Salix* (willow trees). The black cottonwood *Populus trichocarpa* was the first tree to have its genome sequenced and made available to the scientific community (Tuskan *et al*., 2006). Poplar is a model perennial plant widely used to study growth- and immunity-related processes at the molecular and cellular levels (Bradshaw *et al*., 2000; Jansson & Douglas, 2007; Duplessis *et al*., 2009; Hacquard *et al*., 2011). Investigations of the poplar immune system revealed striking differences compared to annual plants; notably, in terms of immune receptor content and diversity, phytohormonal regulation, and defense peptide diversity (Kohler *et al*., 2008; Hacquard *et al*., 2011; Ullah *et al*., 2022).

In 2007, a transcriptomic analysis of poplar leaves revealed an orphan gene called *RUST INDUCED SECRETED PEPTIDE* (*RISP*, hereafter renamed *PtRISP1*) as the most-induced gene during the effective immune response to a rust pathogen infection (Rinaldi *et al*., 2007). PtRISP1 is cationic, thermostable, composed of 60 amino acids in its mature form, and secreted into the apoplast in *Nicotiana benthamiana* (Petre *et al*., 2016). The purified peptide directly inhibits the growth of *Melampsora larici-populina* both *in vitro* and on poplar, and triggers poplar cell culture alkalinization (Petre *et al*., 2016). The *PtRISP1* gene resides next to a *LRR-RP* gene (hereafter named *Populus trichocarpa RISP-ASSOCIATED LRR-RP*; *PtRALR*), and both genes are coregulated in response to biotic or abiotic stress, suggesting a functional link between their products (Petre *et al*., 2014).

The present study aimed at evaluating the diversity and evolution of *RISP* and *RALR* gene families in Salicaceae, and determining whether RALRs recognize RISPs to activate immune signaling. To reach the first objective, we used a phylogenomic approach to inventory and analyze *RISP* and *RALR* genes in Salicaceae. To reach the second objective, we combined protein biochemistry with *in vitro* and *in planta* functional assays to characterize two purified RISPs and to evaluate their ability to trigger immune signaling in a RALR-dependent manner. Overall, this study concludes that *RISP* and *RALR* genes belong to gene families that specifically evolved as clusters in poplar and willow, and that two divergent RISP/RALR pairs from poplar and willow function together to trigger immune signalling.

## Materials and Methods

### Biological material

*N. benthamiana* plants were grown from in-house obtained seeds in soil at 23 °C either in a phytotron chamber for confocal microscopy assays (60 % of relative humidity, and a 16 h photoperiod at 400 µmol.s^-1^.m^-2^) or in a greenhouse for ROS burst and MAPK activation assays. Poplar hybrids (*Populus tremula x Populus alba* clone INRAE 717-1B4) were propagated *in vitro* in test tube from internodes transplant in sterile Murashige and Skoog (MS) medium at pH 5.9-6.0 complemented with 10 ml.l^-1^ of vitamin solution (100 mg.l^-1^ of nicotinic acid, pyridoxine HCl; thiamine, calcium pantothenate; L-Cysteine hydrochloride and 1 ml of biotin solution at 0.1 mg.ml^-1^ in EtOH 95%) in a growth chamber at 23 °C and with a 16-h photoperiod at 50 µmol.s^-1^.m^-2^. The *Escherichia coli* strain BL21 (DE3) psBET and the *Agrobacterium tumefaciens* strain GV3101 (pMP90) were used for the protein production for purification and for the transient protein expression in *N. benthamiana*, respectively. Urediniospores of *Melampsora larici-populina* (isolate 98AG31) were obtained as previously described (Rinaldi *et al*., 2007) and stored as aliquots at -80 °C.

### *In silico* sequence analyses

To identify *RISP* and *RALR* genes in Salicaceae genomes, we searched the predicted proteomes of seven Salicaceae individuals available on Phytozome v13 (https://phytozomenext.jgi.doe.gov/), using the BlastP tool and using the amino acid sequences of PtRISP1 or PtRALR as queries (see Supporting Information Datasets S1, and S2 for details). We also searched the NCBI nr database as well as all available predicted proteomes on the Phytozome portal for additional sequences. The relative positions of *RISP* and *RALR* genes in the genomes were estimated with the JBrowse tool on the Phytozome portal. The sequences of RPL30, RXEG1, Cf9, RPL23, and INR were retrieved from the UniProt and The Arabidopsis Information Resource (TAIR) databases. All the sequences used in this study are archived in Supporting Information Dataset S1 and the Text File S1. Salicaceae-specific LRR-RP sequences were obtained from a previous study (Petre *et al*., 2014) or identified within the predicted proteome of *S. purpurea* on the Phytozome portal.

### Plasmid construction

Binary vectors were built using the Golden Gate modular cloning technology, with cloning kits and protocols described previously (Weber *et al*., 2011; Engler *et al*., 2014; Petre *et al*., 2017). Briefly, coding sequences were obtained by DNA synthesis (Genecust S.A.S, BOYNES, France) or PCR cloning from poplar cDNAs and then sub-cloned into pAGM1287 vectors to create level 0 modules with AATG-TTCG compatible overhangs. The level 0 module was then assembled into a level 1 binary vector (pICH47742 or vector from the same series), along with a short version of the 35S promoter (pICH51277; GGAG-AATG compatible overhangs), the coding sequence of a C-terminal tag such as a mCherry (pICSL50004; AATG-GCTT compatible overhangs) or a GREEN FLUORESCENT PROTEIN (GFP) (pICSL50008; AATG-GCTT compatible overhangs), and a combined 3’ UTR / OCS terminator (pICH41432; GCTT-CGCT compatible overhangs) (Dataset S1; Fig. S1a). The coding sequence of P19 suppressor of gene silencing (pICH44022, AATG-GCTT compatible overhangs) was assembled into a level 1 binary vector (pICH47761) along with a short version of the 35S promoter and a combined 3’ UTR / OCS terminator (Jay *et al*., 2023). Multigene (i.e., level 2) vectors were built by combining DNA fragments from appropriate and compatible level 1 vectors (Fig. S2). Vectors for bacterial protein expression were build using the restriction/ligation technology and a collection of pET vectors as previously described (Petre *et al*., 2016). Briefly, coding sequence of the mature (without signal peptide) form of SpRISP1 was obtained by DNA synthesis directly cloned into pET15b vector (insert between NdeI/BamH1 restriction sites), with an N-terminal hexahistidine tag encoded by the vector replacing the predicted signal peptide (Genecust S.A.S, BOYNES, France) (Fig. S1a). All purified plasmids were stored in double distilled water (ddHOH) at -20 °C until further use. All vectors obtained and used in this study are listed in Dataset S1, along with the amino acid sequence of relevant proteins.

### Phylogenetic analyses

To build phylogenetic trees, we used a five-step pipeline. Firstly, we performed an amino-acid alignment using the muscle algorithm implemented in the Seaview software (Gouy *et al*., 2010). Secondly, we used this alignment to identify the best substitution model with the IQTREE web server (Trifinopoulos *et al*., 2016). Thirdly, we selected suitable positions to build a tree in the alignment of the first step with Gblock algorithm (for RISPs) or selected the positions matching the C3-D domain (for LRR-RPs). Fourthly, we built a maximum likelihood tree (PhyML tool, with model identified in step 2, 100 to 1000 bootstraps, and default parameters) and archived unrooted trees as text files (Supporting Information text file S2). Finally, we used the graphical software Figtree (http://tree.bio.ed.ac.uk/software/figtree/) as well as Microsoft PowerPoint to analyse and render final trees displayed in the manuscript.

### Protein expression in *E. coli* and purification by affinity chromatography

To express proteins in the cytosol of *E. coli*, we inserted pET15b vectors into *E. coli* strains BL21 (DE3) psBET and selected transformants on solid LB broth with appropriate antibiotics at 37°C. Protein expression was induced during the exponential phase of growth of the bacteria by adding isopropyl β-D-thiogalactopyranoside (IPTG) to the bacterial culture at a final concentration of 100 μM for 4 h at 37 °C. The bacteria cultures were then centrifuged for 15 min at 5000 rpm at 4 °C. Pellets were resuspended in 15 mL of TE-NaCl buffer (50 mM Tris-HCl pH 8.0, 1 mM Ethylenediaminetetraacetic acid (EDTA), 100 mM NaCl) and stored at -20 °C. To purify the histidine-tagged proteins, the pellets were sonicated, and the soluble and insoluble fractions were separated by centrifugation for 30 min at 20,000 rpm at 4 °C. The soluble fractions were loaded onto an immobilized-metal affinity chromatography (IMAC) column (Ni Sepharose™ 6 Fast Flow, Cytiva, Sweden) using a peristaltic pump. Successive washing steps were performed with a washing buffer (50 mM Tris-HCl pH 8.0, 300 mM NaCl, 20 mM imidazole) to remove contaminants. Proteins were eluted using 20 mL of elution buffer (50 mM Tris-HCl pH 8.0, 300 mM NaCl, 250 mM imidazole) and concentrated to 1-2 mL by ultrafiltration using a Vivaspin® Turbo Centrifugal Concentrator (Sartorius, United Kingdom). To remove imidazole, proteins were transferred into a dialysis membrane (Standard RC Tubing, MWCO: 6-8 kDa, diameter 6.4 mm, Spectrum Laboratories, Inc) and incubated under agitation in dialysis buffer (30 mM Tris-HCl pH 8.0, 200 mM NaCl, 1 mM EDTA) overnight and two additional hours in a new dialysis solution. Protein concentration was determined by a spectrophotometric analysis of protein absorbance at 280 nm and protein integrity was estimated by 15 % SDS-PAGE/CCB staining (Fig. S1b). Purified proteins were stored at 4 °C and used within 30 days.

### Thermo-stability, spore pull down, and inhibition of spore germination assays

Protein thermostability, spore pull-down and *in vitro* inhibition of spore germination assays were performed as described previously in Petre *et al*., 2016. Briefly, to perform the protein thermostability assay, purified proteins were incubated 10 min at 95 °C and centrifuged.

Supernatant was collected and proteins were visualized on SDS-PAGE 15 % acrylamide followed by Coomassie blue staining. Purified PtRISP1 and GFP were used in these bioassays as positive and negative controls, respectively.

### Transient protein expression in *N. benthamiana*, laser-scanning confocal microscopy assays, and image analysis

Transient protein expression and confocal microscopy were performed as previously described (Petre *et al*., 2017). Briefly, binary vectors were inserted into *A. tumefaciens* strain GV3101 (pMP90); the bacteria carrying the vector were then infiltrated in the leaves of three to five-week-old *N. benthamiana* plants. Microscopy analyses were performed with a ZEISS LSM 780 (Zeiss International) laser-scanning confocal microscope with the 40x objective using the methods for image acquisition and interpretation described in Petre *et al*., 2021. The fluorescence of GFP, mCherry, and chlorophyll were observed using the following excitation/emission wavelengths: 488/500-525 nm, 561/580-620 nm, and 488/680-700 nm, respectively. The fluorescence intensity was measured by using the “Measure” tool on Fiji software (https://fiji.sc/), and data were exported in a spreadsheet and analyzed with Microsoft Excel.

### MAPK activation assays

Leaf disks of *N. benthamiana* transiently expressing RALR and PtSOBIR1 were harvested 3 days-post-infiltration with a biopsy punch (8-mm diameter). Eight leaf disks were infiltrated with 1 µM flg22 or 100 µM of purified PtRISP1/SpRISP1 and incubated for 0, 15, or 30 min prior to flash-freezing and grinding in liquid nitrogen. Proteins were extracted by incubation of the leaf powder in Laemmli buffer (62.5 mM Tris-HCl pH 6.8, 8.3% glycerol, 2% SDS, 0.017% bromophenol blue) with 100 mM dithiothreitol (DTT) for 10 min at 95 °C. The samples were centrifuged at 13,000 rpm and the proteins were separated by SDS-PAGE. Proteins were then transferred onto a polyvinylidene fluoride (PVDF) membrane, which was then incubated with the primary antibody α-pMAPK (Phospho-p44/42 MAPK (erk1/2) 1:5,000; Cell Signalling technology) and the secondary antibody (Sigma anti-Rabbit 1:10,000). Antibody signals onto membranes were imaged with a ChemiDoc Imaging System (Bio-Rad). Coomassie brilliant blue (CBB) staining of the proteins onto the membranes was used as a loading control.

### ROS burst assay

Leaf disks of *N. benthamiana* transiently expressing RALR and PtSOBIR1 were harvested 3 days-post-infiltration with a biopsy punch (4-mm diameter). Disks were placed in a 96-well plate, incubated in 100 µL distilled water, and kept at room temperature overnight. Prior the ROS quantification, water was replaced with 100 µL of assay solution (0.5 µM L-012, 10 µg.ml^1^ horseradish peroxidase [HRP], 100 nM flg22 in water or 100 µM of purified RISP, or TE buffer pH 8) and light emission was measured immediately with a Spark microplate reader (Tecan, Switzerland). Relative light units (RLUs) were collected in 60 s intervals for 60- or 90-min. Data were exported in a spreadsheet and analyzed with Microsoft Excel.

### Stomatal closure assay

Stomatal closure assay was performed with leaves of three to five-month-old plants of *P. tremula x P. alba* clone INRAE 717-1B4 grown *in vitro*. Full leaves were harvested and incubated in a stomata opening buffer (10 mM MES, 50 mM KCl, pH 6.15) (Shen *et al*., 2021) for 2 h under light (50 µmol.m^-2^.s^-1^). Leaves were then incubated for 2 h in purified PtRISP1 or GFP to a final concentration of 100 µM, in water under light condition (open stomata control), or in water under dark condition (closed stomata control). Leaves were then mounted in water between a glass slide and a cover slip and images of randomly selected positions of the abaxial side of the leaves were recorded with a light microscope with a 40x water immersion objective and a camera (Lordil). The stomatal opening widths and lengths were measured on images using Fiji and the width/length ratio was calculated to evaluate the stomatal aperture (raw data are available in Supporting Information Dataset S4). Statistical analyses were performed on RStudio using Wilcoxon rank sum test with continuity correction.

## Results

### Clusters of *RISP* and *RALR* genes evolved specifically in Salicaceae

To determine if PtRISP1 belongs to a gene family, we comprehensively searched for PtRISP1 homologs in publicly available predicted proteomes. In total, the search identified 24 such homologs (hereafter RISPs) in 8 different genomes of 7 Salicaceae species (Dataset S2). Those 24 *RISP* genes group among 8 clusters of 2 to 4 genes harbored by chromosome 9 (one cluster per genome); except for *P. trichocarpa* which presents two clusters (a second cluster being present on the small scaffold 502). The 24 RISP family members vary in size from 76 to 83 amino acids (50 to 58 amino acids in their mature form) and exhibit an average percentage identity of 68 % (Fig. **1a**). We found no RISP outside poplar or willow, suggesting that the RISP family evolved specifically in Salicaceae species. The phylogenetic analysis shows that poplar and willow RISPs group into two well-supported phylogenetic clades, suggesting that the family evolved from a single ancestral gene that emerged in the ancestor species of poplars and willows approx. 60 million years ago (Fig. **1a**) (Liu *et al*., 2022). RISPs predicted signal peptides are highly conserved (mean p-distance of 0.158 ± 0.11), whereas RISPs mature forms differ more (mean p-distance of 0.458 ± 0.18). Despite this sequence variability, RISPs mature forms present 4 regions with noticeable and conserved properties: i) a N-terminal region with a predicted alpha-helical structure, ii) a hydrophilic region, iii) a positively charged region (average net charge of positive 6 ± 1.6) and iv) a C-terminal negatively-charged region (average net charge of negative 2.7 ± 0.8) (Fig. **1a**; Dataset S2). Also, RISPs have four fully conserved cysteines in their mature form and present high predicted isoelectric points (average of 9.4 ± 0.3) (Fig. **1a**). Altogether, these results suggest that RISP evolved as clusters, specifically and recently in Salicaceae species to form a diverse family of cationic secreted peptides.

**Figure 1.**
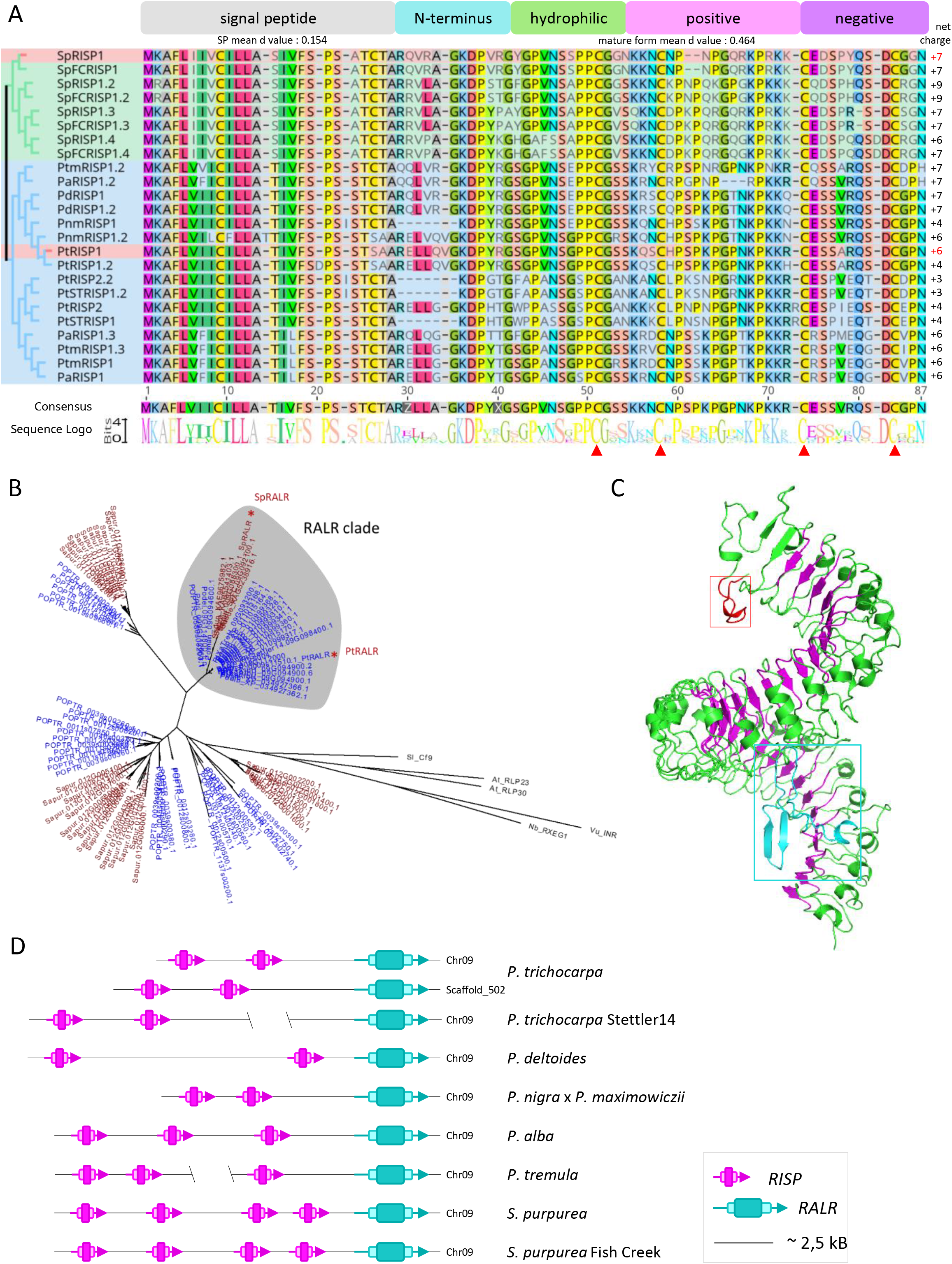
Clusters comprising RISP and RALR genes evolved specifically in Salicaceae. (A) Alignment of 24 RISP amino acid sequences identified in the *Salicaceae* genomes. The alignment matches the phylogenetic tree represented on the left side: *Populus* and *Salix* sequences are highlighted in blue and green, respectively (Supporting Information Dataset S2). The two RISPs investigated in this study are highlighted in red. At the top of the alignment, five regions with various properties were identified: a predicted signal peptide (grey box), a N-terminus region (blue box), a hydrophilic region (green box), a positively charged region (pink box) and a negatively charged region (purple box). At the bottom of the alignment, the consensus sequence and the web logo are represented; the red arrowheads point the four highly conserved cysteines. Net charge for each peptide is indicated (column on the right). (B) Phylogenetic tree generated from *Populus* (blue) and *Salix* (red) LRR-RPs as well as selected LRR-RPs from model plants (*Arabidopsis thaliana*; *Nicotiana benthamiana; Solanum lycopersicum*; *Vigna unguiculata*) (black). All RALRs gather into a well-separated clade (RALR clade highlighted in grey). Within the RALR clade, the willow sequences gather into a separate sub-clade. Red dots point to SpRALR and PtRALR. (C) AlphaFold2-predicted tridimensional structure of *Pt*RALR, with the leucine-rich repeat region in green and the LRR motifs in pink (pLDDT=83,1). The red and blue squares indicate the N-loopout (red) and the Island domain (cyan), respectively. Movie and additional 3D structures are available in Fig. S4 (Jumper *et al*., 2021). (D) Schematic representation of the clusters of *RISP* (magenta) and *RALR (cyan)* genes on the chromosomes or scaffolds in the genomes of *P. trichocarpa, P. trichocarpa* Stettler14, *P. deltoïdes, P. nigra* x *P. maximowiczii, P. tremula* x *P. alba (tremula or alba haplotypes), S. purpurea,* and *S. purpurea* Fish Creek. Raw data is available in Supporting Information Dataset S3.

To evaluate how *PtRALR* evolved within the *LRR-RP* gene family, we comprehensively searched for PtRALR homologs in publicly available predicted proteomes as well as in the NCBI protein database. In total, this search identified only 25 such homologs (hereafter RALRs) belonging to ten different Salicaceae species (Text File S1). The 25 RALRs display an average amino acid similarity of 88.5 % (ranging from 70 % to 99.5 %) and an average length of 1060 amino acids (ranging from 1045 to 1075; PtRALR comprising 1046 amino acids). All RALRs gather into a well-separated clade (hereafter the RALR clade) within a phylogenetic tree of LRR-RPs; the RALR clade itself residing within a large clade of Salicaceae LRR-RPs. Within the RALR clade, the willow sequences gather into a separate sub-clade (Fig. **1b**). Interestingly, among the 25 RALRs, 8 originate from the 8 Salicaceae genomes present on the Phytozome portal; those 8 *RALR* genes all reside within the *RISP* clusters, immediately downstream of the *RISP* genes (Fig. **1d**; Dataset S3). Thus, all *RISP* and *RALR* genes identified in the available Salicaceae genomes so far cluster together, in such a way that the clusters comprise one *RALR* gene and two to four *RISP* genes. Overall, these findings suggest that clusters comprising *RISP* and *RALR* genes evolved and diversified from a common ancestor cluster in Salicaceae species.

We hypothesized that the products of *RISP* and *RALR* genes present in the same cluster function together to trigger immune signaling. To functionally test this hypothesis, we selected two pairs of clustered *RISP* and *RALR* genes: *PtRISP1*/*PtRALR* in poplar, and *SpRISP1*/*SpRALR* in willow (Fig. **1d**; Fig. S3). Both pairs encode divergent proteins, as the mature forms of PtRISP1 and SpRISP1 as well as PtRALR and SpRALR exhibit only 59 % and 85 % of amino acid identity, respectively (Fig. S3). PtRALR and SpRALR present all the canonical domains of LRR-RPs: a predicted N-terminal signal peptide, a cysteine-rich domain, a leucinerich repeat region with 33 LRRs, an acid-rich domain, a transmembrane helix, and a cytosolic tail (Fig. **1c**; Figs. S4; S5). The AlphaFold2-generated tridimensional models of both PtRALR and SpRALR predict the canonical superhelix fold of the LRR domain, that comprises the N-terminal loop (N-loopout) and C-terminal island domain (ID) involved in ligand recognition in other LRRRPs (Fig. **1c**; Figs. S4; S5) (Matsushima & Miyashita, 2012; Sun *et al*., 2022; Snoeck *et al*., 2023).

### SpRISP1 and PtRISP1 exhibit similar biophysical properties and antimicrobial activities

A previous study showed that PtRISP1 accumulates in the apoplast in *N. benthamiana*, is thermostable, and binds and inhibits the germination of urediniospores of *M. larici-populina* (Petre *et al*., 2016). We aimed at determining whether SpRISP1 presents similar biophysical properties and antimicrobial activities. To this end, we first transiently expressed SpRISP1mCherry fusion in *N. benthamiana* leaves and determined its accumulation pattern by laser scanning confocal microscopy. This analysis showed that SpRISP1 and PtRISP1 (used as a positive control) exclusively accumulate in the apoplast without overlapping with the signal of a free GFP (used as a nucleo-cytoplasmic marker) (Fig. **2a**). In addition, western blot analyses revealed the presence of RISP-mCherry fusions and SP-Ramya3A-mCherry (apoplastic control) in apoplastic fluids from *N. benthamiana* leaves, whereas intracellular GFP was only detected in total leaf protein extracts (Fig. S6). Then, we obtained the mature form of SpRISP1 as a purified protein produced in *E. coli* and observed that the protein remains soluble after heat treatment for 10 min at 95 °C, similar to PtRISP1 (Fig. **2b**). Protein-spore pull-down assays showed that SpRISP1 attaches to urediniospores, similar to PtRISP1 (positive control), whereas a GFP negative control did not (Fig. **2c**). Finally, inhibition of germination assays revealed that a solution of 100 µM SpRISP1 inhibited the germination of *M. larici-populina* urediniospores, similar to a PtRISP1 positive control (germination rates of 22 % ± 7 and 8.5 % ± 6 respectively), as opposed to the mock treatment which had a high germination rate of over 85 % ± 8 (Fig. **2d**). Altogether, these results indicate that SpRISP1 accumulates in the apoplast, is thermostable, and interacts with *M. larici-populina* urediniospores and inhibits their germination. As PtRISP1 and SpRISP1 belong to the two major and divergent sub-clades of their family, these findings suggest that RISP family members retained similar biophysical properties and activities throughout evolution.

**Figure 2.**
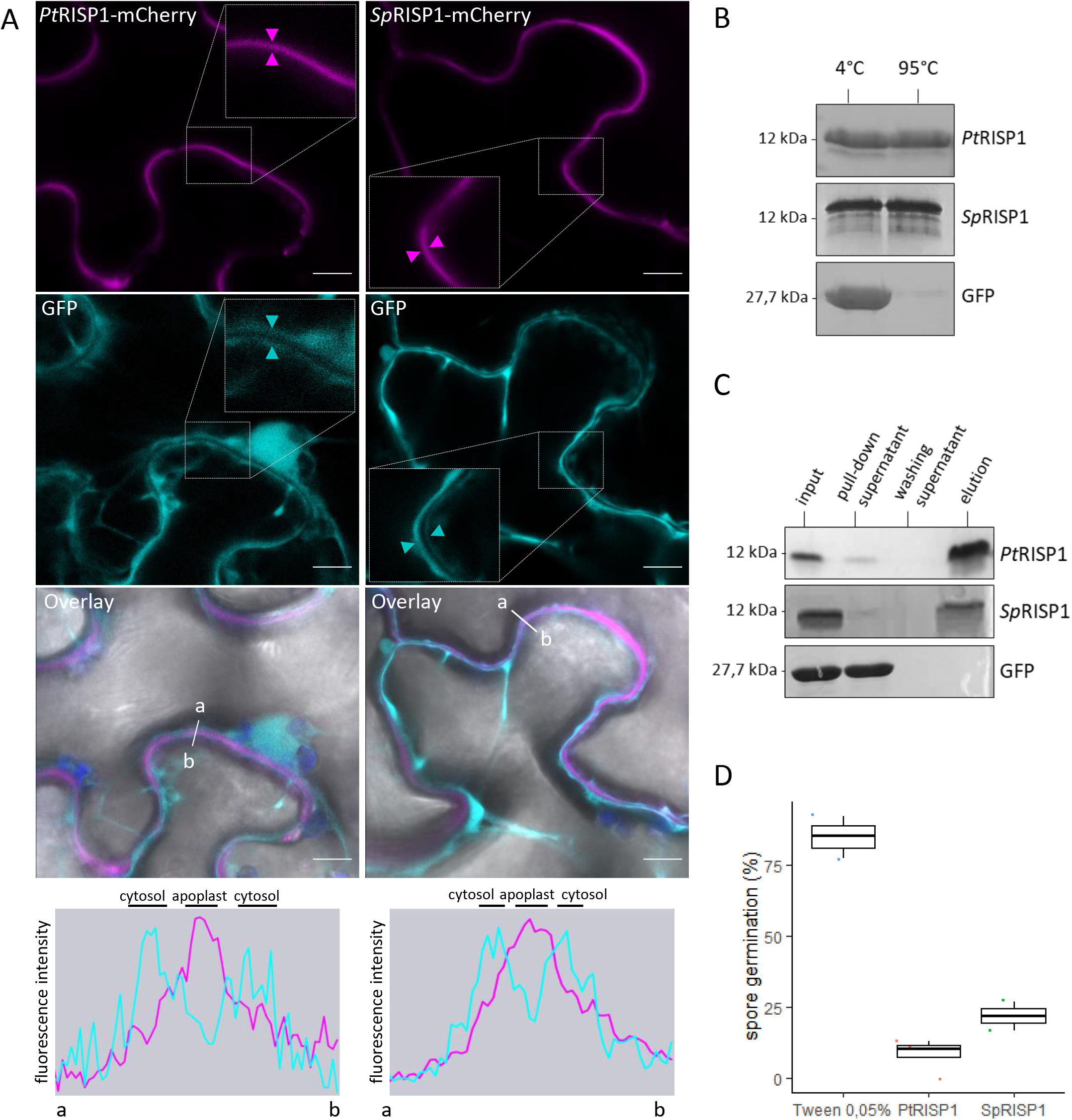
SpRISP1 and PtRISP1 show similar biophysical properties and antimicrobial activities. (A) PtRISP1 and SpRISP1 accumulate in the apoplast. Confocal microscopy images of *N. benthamiana* leaf epidermal cells transiently co-accumulating SpRISP1-mCherry or PtRISP1-mCherry (in cyan) with a free GFP (in magenta) used as a nucleo-cytoplasmic marker, acquired through three independent agroinfiltration assays. Scale bars = 10 µm. Image inserts show 2x zoomed areas, the magenta arrowhead indicates the apoplast; the cyan arrowheads indicate cytosols. Fluorescence intensity graphs show mCherry (magenta) and GFP (cyan) signals measured along the white line from a to b. Detection of both RISP-mCherry fusions in the apoplastic fluid from *N. benthamiana* leaves is show in Supporting Information Fig. S6. (B) SpRISP1 and PtRISP1 are thermosoluble. Purified PtRISP1, SpRISP1, and GFP proteins were incubated at 95 °C for 10 min; soluble proteins before (4 °C) or after heating (95 °C) were visualized by SDS-PAGE/CBB staining. (C) SpRISP1 and PtRISP1 interact with *Melampsora laricipopulina* urediniospores *in vitro*. Purified PtRISP1, SpRISP1, and GFP proteins were incubated with urediniospores. Purified proteins of the different fractions collected were visualized by SDS-PAGE/CBB staining. Input: protein solution before centrifugation. Pull-down supernatant: supernatant after the first centrifugation. Washing supernatant: supernatant after the second centrifugation for washing. Elution: supernatant after incubation at 95°C in Laemmli buffer. (D) PtRISP1 and SpRISP1 inhibit *M. larici-populina* urediniospore germination. *In vitro* inhibition of germination assays were performed on water-agar medium with 100 µM of purified PtRISP1 or SpRISP1 boiled 10 min at 95 °C, or with Tween 0.05 % (mock treatment). The percentage of germination was calculated 6h after the first contact between the spores and the proteins.

### PtRALR and SpRALR accumulate at the plasma membrane in a PtSOBIR1-dependent manner in *N. benthamiana*

To functionally investigate the ability of RALRs to recognize RISPs, we used transient expression assays in *N. benthamiana* (as Salicaceae species are limitedly amenable to reverse genetics). Firstly, we aimed at accumulating RALR-GFP fusions in leaf cells to characterize their subcellular localization, by co-expressing PtRALR-GFP or SpRALR-GFP fusions with PtRISP1mCherry (used as an apoplastic marker) and the P19 protein (a silencing suppressor) in leaves by agroinfiltration. This assay revealed a weak GFP signal at the cell periphery, which did not overlap with the mCherry signal (Fig. **3**). Of note, the accumulation of RALR-GFP fusions required the presence of the P19 protein, as we observed no fluorescent signal in assays without P19. These first results indicate that RALR-GFP fusions can accumulate in *N. benthamiana*, but at low levels, which precludes further functional analyses.

**Figure 3.**
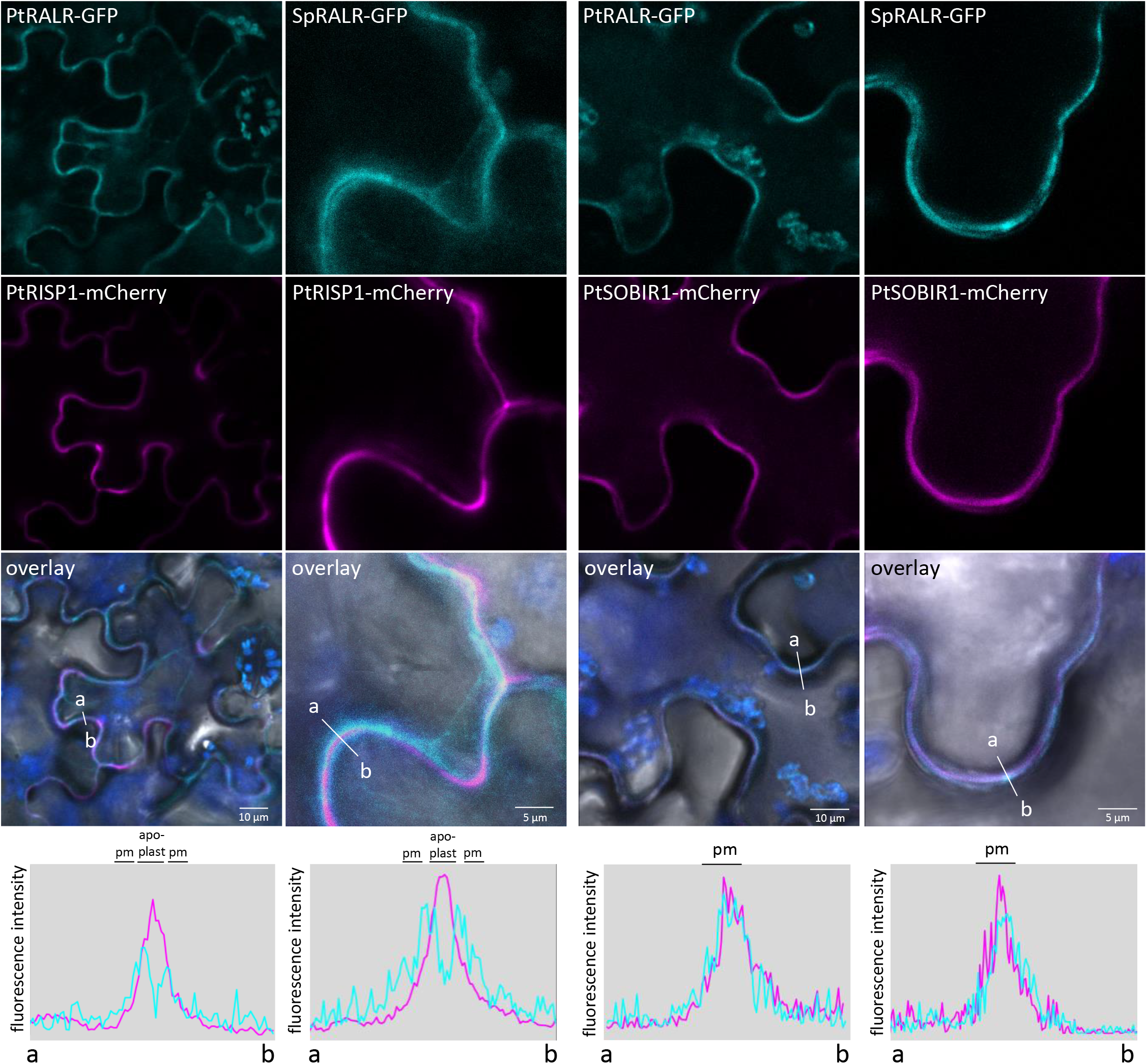
PtRALR and SpRALR accumulate at the plasma membrane in a PtSOBIR1-dependent manner in *N. benthamiana*. Confocal microscopy images of *N. benthamiana* leaf epidermal cells transiently co-accumulating PtRALR-GFP and SpRALR-GFP (cyan images on top), PtRISP1mCherry as apoplastic marker or PtSOBIR1-mCherry as plasma membrane marker (magenta images in the middle) and P19 (suppressor of gene silencing) protein. Live cell imaging was performed with a confocal microscope three days after infiltration. The overlay images combine the GFP, mcherry, chlorophyll (blue) and bright field. Scale bars=5µm or 10 µm. Fluorescence intensity graphs show mCherry (magenta) and GFP (cyan) signals measured along the white line from a to b.

The presence of the adaptor kinase SOBIR1 was shown to assist the accumulation of LRR-RPs in plant cells (Liebrand *et al*., 2013, 2014; Böhm *et al*., 2014). To improve our ability to express RALR-GFP fusions in *N. benthamiana*, we aimed at co-expressing them with a homolog of SOBIR1 from a Salicaceae species. To this end, we cloned the coding sequence of one of the two SOBIR1 homologs in *P. trichocarpa* (hereafter PtSOBIR1). PtSOBIR1 shares a 64 % amino acid identity with *A. thaliana* SOBIR1, and comprises both a conserved C-terminus and a GXXXG dimerization motif in the transmembrane domain (Bi *et al*., 2016) (Fig. S7). As PtSOBIR1 shares a high amino acid identity (88 %) with its closest homolog in *Salix*, we used PtSOBIR1 for the assays with SpRALR. As anticipated, PtSOBIR1-mCherry fusions clearly and specifically accumulated at the plasma membrane in *N. benthamiana* (Fig. S8a). Co-expression of RALRGFP, P19, and PtSOBIR1-mCherry fusions revealed a well-detectable co-accumulation of fluorescent signals at the plasma membrane (Fig. **3**). Western blot analyses revealed the presence of intact PtRALR-GFP, SpRALR-GFP, and PtSOBIR1-mCherry in leaf protein extracts (Fig. S8b). Notably, PtSOBIR1 promoted the accumulation of RALR-GFP fusions at the plasma membrane, as we could observe a GFP signal in the absence of P19. In conclusion, these results indicate that both PtRALR and SpRALR can accumulate at the plasma membrane in *N. benthamiana*, and that the presence of PtSOBIR1 facilitates this accumulation. As no cell death or leaf stress symptoms were detectable, we surmised that transient assays would be suitable to study RALR-mediated immune signaling activation.

### Purified RISPs trigger immune signaling in a RALR-dependent manner in *N. benthamiana*

To test whether RALRs are sufficient to confer RISP-responsiveness to *N. benthamiana*, we combined transient assays with purified peptide treatments followed by the dynamic quantification of ROS and phosphorylated MAPKs. On the one hand, an exogenous treatment with purified PtRISP1 of leaf disks expressing PtRALR and PtSOBIR1 triggered a ROS burst that peaked at 30 min, as well as a strong accumulation of phosphorylated MAPKs 15 and 30 min post-treatment; with intensities comparable to those triggered by the flg22 positive control (Fig. **4a, b**). On the other hand, the same experiment performed with the SpRISP1/SpRALR pair revealed weaker ROS and phosphorylated MAPKs accumulation, although both showed transient accumulation patterns. Altogether, we conclude that the co-expression of RALRs and PtSOBIR1 in *N. benthamiana* leaves confers RISP-responsiveness, suggesting that PtRALR and SpRALR recognize PtRISP1 and SpRISP1, respectively, and that this recognition rapidly initiates immune signaling events.

**Figure 4.**
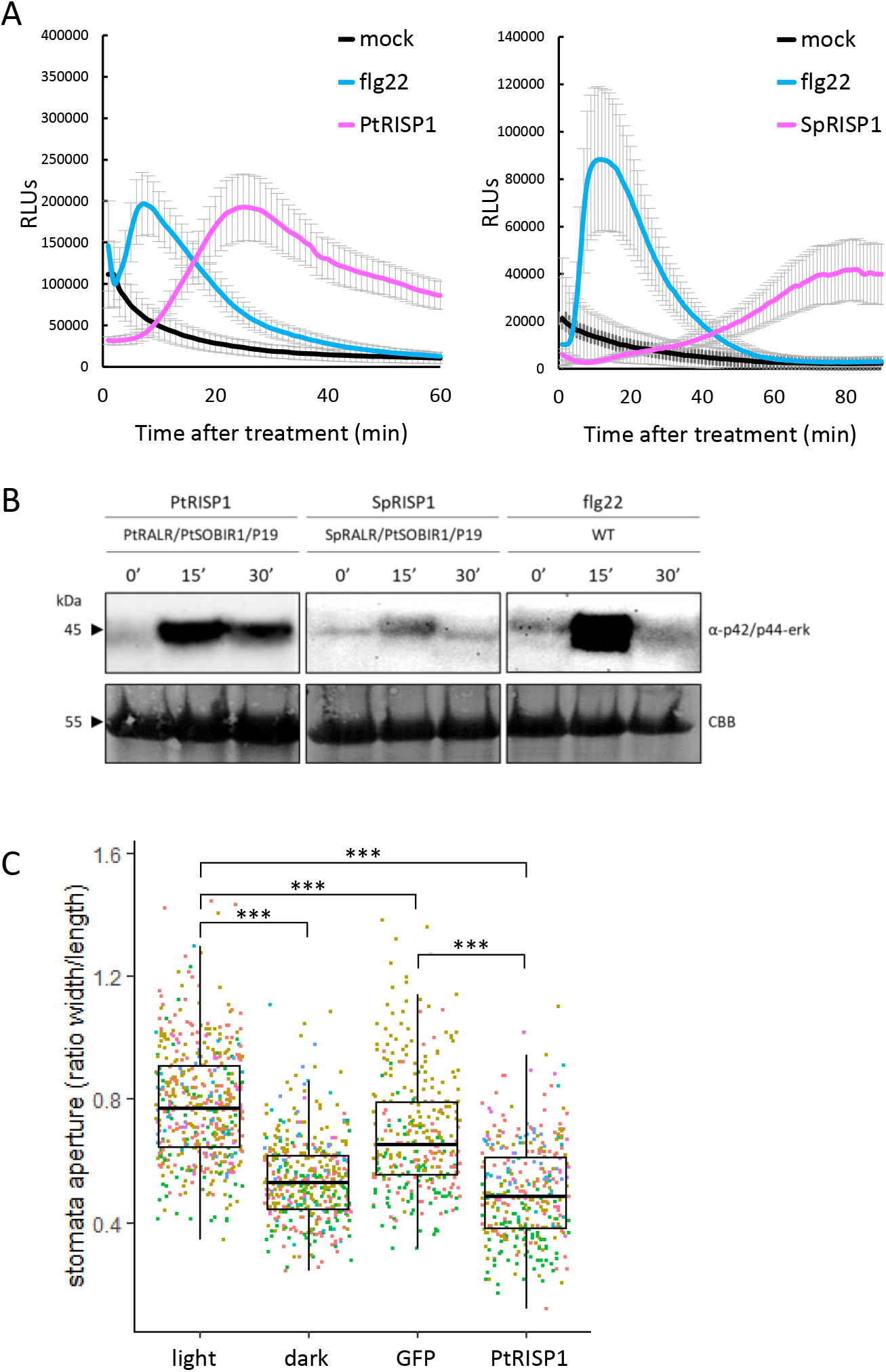
RISPs induce immune signaling in RALR-expressing *N. benthamiana* or poplar. (A) ROS production in leaf disks expressing PtRALR, PtSOBIR1 and P19 proteins in *N. benthamiana* plants treated with 100 μM PtRISP1 (left graph - magenta) or 100 μM SpRISP1 application (right graph - magenta), 1 µM flg22 (cyan), and TE buffer (mock treatment) (black). Error bars represent standard errors (B) Western blot using α-p42/p44-ERK recognizing phosphorylated MAPKs in *N. benthamiana* leaf disks expressing PtRALR/PtSOBIR1/P19 or SpRALR/PtSOBIR1/P19 or WT control and treated with 100 µM of PtRISP1, 100 µM of SpRISP1, or 1 µM of flg22, respectively, for 0, 15, and 30 min. Membranes were stained with CBB as loading control. (C) PtRISP1 induces stomatal closure in *P. tremula x P. alba* leaves (cultivar 717-1B4) from culture *in vitro.* Poplar leaves were incubated 2 h in water or water containing 100 µM of purified GFP, PtRISP1 under light or placed in darkness. Images of abaxial side of leaves were taken with a light microscope and the ratio width/length of stomata was measured using ImageJ. The different colors of the dots correspond to the six independent replicates. Asterisks stand for p-value < 0.05 from Pairwise comparisons using Wilcoxon rank sum test with continuity correction (n=6; 1574 stomata analyzed).

### Purified PtRISP1 triggers stomatal closure in poplar

To determine if PtRISP1 can activate immune responses in its organism of origin, we established a stomatal closure assay in poplar. Briefly, we treated detached leaves of *in vitro*grown hybrid poplars with purified RISP proteins for two hours, then estimated the stomatal aperture by using the width/length ratio method (Thor *et al*., 2020). This assay showed that PtRISP1 treatment reduces stomatal aperture (ratio of 0.50 ± 0.16) similarly to the dark positive control (ratio of 0.54 ± 0.13), whereas leaves incubated with purified GFP or a mock treatment showed higher stomatal aperture (ratio of 0.69 ± 0.2 and 0.78 ± 0.19, respectively) (Fig. **4c**). Statistical analyses indicated that stomatal aperture was significantly different between PtRISP1, light condition, and GFP, whereas no significant difference was observed between PtRISP1 and the dark condition used as a positive control. Thus, exogenous treatment of PtRISP1 triggers a rapid and strong stomatal closure, suggesting that PtRISP1 is sufficient to elicit an immune-related response in poplar leaves.

## Discussion

This study reports that clusters of *RISP* and *RALR* genes evolved recently and specifically in Salicaceae species, and that RISP family members function as defense peptides with both antifungal and elicitor activities; the elicitor activity being mediated by their clustered RALR. This section discusses the multi-functionality of plant defense peptides, the surprising clustering of ligand/receptor pairs in plants, the efforts required for the characterization of LRR-RPs in non-model species, and the original position of RALRs as LRR-RPs recognizing phytocytokines.

### The characterization of plant functional analogs of metazoan host-defense peptides emerges as research front

We showed that RISP family members simultaneously exhibit antimicrobial and immunomodulatory activities, making RISPs functional analogs of metazoan host-defense peptides (HDPs). The characterization of HDPs analogs in plants is emerging as a research front (Petre, 2020; Han *et al*., 2023). Notably, recent studies have reported two superfamilies of plant defense peptides, namely PATHOGENESIS-RELATED PROTEIN 1 (PR1) and SERINE RICH ENDOGENOUS PEPTIDES (SCOOPs), with members having antimicrobial and immunomodulatory activities (Neukermans *et al*., 2015; Yu *et al*., 2020; Guillou *et al*., 2022; Han *et al*., 2023). PR1 superfamily members are well-characterized inhibitors of microbial growth (Niderman *et al*., 1995), and PR1 is cleaved to release the C-terminal CAP-derived peptides (CAPEs) that activates plant immune responses (Chen *et al*., 2014, 2023; Sung *et al*., 2021). PR1 superfamily members are also targeted by pathogen effectors that prevent CAPE1 cleavage, demonstrating the importance of this process in plant immunity (Lu *et al*., 2014; Sung *et al*., 2021). Furthermore, the divergent superfamily of SCOOPs comprises members exhibiting antifungal activities (Neukermans *et al*., 2015; Yu *et al*., 2020). SCOOP peptides are recognized by the *A. thaliana* LRR-RK MALE DISCOVERER 1-INTERACTING RECEPTOR LIKE KINASE 2 (MIK2) to induce immunity (Hou *et al*., 2021; Rhodes *et al*., 2021; Zhang *et al*., 2022) In addition to these two well-studied families, defensin and thaumatin-like protein families may also harbor functional analogs of HDPs (Petre, 2020). Future studies on plant multifunctional immune peptides may help reveal their valorization potential in agriculture as versatile ’Swiss-army knife’ molecules (Hilchie *et al*., 2013; Sun *et al*., 2023). Such studies could also improve our understanding of the eukaryotic immune systems, for instance by highlighting how metazoans and plants evolved functionally analogous defense peptides.

### Can gene clustering analyses help to identify ligand/receptor pair candidates?

A pilot study screened the poplar genome to reveal that *PtRISP1* and *PtRALR* genes cluster together, and hypothesized a functional link between the two (Petre *et al*., 2014). In this follow-up study, we showed that *RISP* and *RALR* gene family members systematically cluster, and that at least two RISP/RALR pairs function together to trigger immune signaling. The direct clustering of genes encoding phytocytokines and cell-surface receptors is not common. For instance, in *A. thaliana*, the genes encoding well-characterized ligand/receptor pairs, such as PEP1/PEPR1, PIP1/RLK7, or SCOOP12/MIK2, reside on different chromosomes. Indeed, *PEP1, PIP1* and *SCOOP12* genes are located on chromosomes 2, 3 and 5 respectively, whereas the genes encoding their receptors are all located on chromosome 1 in the *A. thaliana* genome (Dataset S3) (Rzemieniewski & Stegmann, 2022). Identifying ligand/receptor pairs is a central goal in both plant or animal biology; though such an endeavor requires experimentally demanding and time-consuming screening approaches (Ramilowski *et al*., 2015; Boutrot & Zipfel, 2017; Siepe *et al*., 2022). Screening genomes for physically associated and co-regulated genes encoding cell-surface receptors and small secreted proteins may help accelerate the identification of ligand/receptor pair candidates.

### Characterizing LRR-RPs from non-model species requires significant efforts to gain genomewide family knowledge, produce molecular material, and implement methodologies

Our study initiated the characterization of LRR-RPs in two species of the Salicaceae family. The 19 LRR-RPs characterized as immune receptors so far belong to only three plant families: *Solanaceae* (12), *Brassicaceae* (6), and *Fabaceae* (1) (Snoeck *et al*., 2023). Though poplar and willow are considered as model trees, they lack strong reverse genetic tools and remain underinvestigated compared with annual models (Marks *et al*., 2023). For instance, rapid transient expression assays are still lacking and cannot be systematically applied. Characterizing LRRRPs from non-model botanical families faces the double challenge of building family-specific background knowledge and acquiring material and methodologies. We tackled the first challenge by performing a comprehensive phylogenomic analysis of LRR-RPs and structural predictions; some of this effort was included in a pilot study that was instrumental to generate hypotheses and select protein candidates (Petre *et al*., 2014). We tackled the second challenge by both using heterologous systems (notably *N. benthamiana* to express RALRs *in planta* and *E. coli* to produce RISPs) and by implementing novel methodologies and protocols (notably the stomatal closure assay in poplar); such efforts and approaches being commonly required in non-models (Petre *et al*., 2016; Lorrain *et al*., 2018). The implementation of methodologies and protocols is time-consuming and often ineffective. For instance, in this study, attempts to implement several bioassays in poplar were unsuccessful while consuming significant human and financial resources (i.e., gene expression induction, cell culture-based assays, seedling growth inhibition assays, protein-protein interaction assays). The molecular resources and methodologies we built here will hopefully facilitate future studies addressing either LRR-RPs and/or the molecular physiology of Salicaceae.

### RALR is the first LRR-RP reported to recognize a phytocytokine

We showed that RALRs mediate the recognition of the phytocytokines, RISPs. To our knowledge, RALR is the first LRR-RP reported to recognize a phytocytokine. Indeed, known receptors of phytocytokines all belong to the superfamily of RKs (Ngou *et al*., 2022; Rzemieniewski & Stegmann, 2022). Among the 19 characterized LRR-RP immune receptors, all but one recognize pathogen-derived peptides (Snoeck *et al*., 2023). Indeed, only INR recognizes a self-molecule; a plant-derived peptide proteolytically generated from a chloroplastic ATP synthase upon caterpillar chewing, which does not qualify as a phytocytokine *per se* (Steinbrenner *et al*., 2020; Rzemieniewski & Stegmann, 2022). The RISP/RALR pairs may serve as models to dissect the LRR-RP-mediated recognition of phytocytokines as well as downstream signaling events and responses. To this end, the functionality of RALR in *N. benthamiana* will be instrumental, along with i) the ability to easily biosynthesize and purify RISPs, ii) the diversity of RISP and RALR sequences we identified to assist structure/function approaches, and iii) the ability to predict *in silico* the structure of RALRs super-helical LRR domain (Fig. **1**; Fig. S4). PtRISP1 was previously shown to undergo processing at its C-terminus by plant proteins (Petre *et al*., 2016); one challenge to further dissect RISP/RALR functioning will also be to identify the exact sequence of that C-terminal peptide.

## Supporting information

DatasetsS1-S4

## Acknowledgements

The authors acknowledge members of the UMR IAM, and notably N. Rouhier, M. MorelRouhier, C. Veneault-Fourrey, F. Lauve-Zannini, and C.P.D. Louet for fruitful discussions, C. Teissier for administrative support, M-L. Ancel, A. Deveau, T. Dhalleine, D. Culot-Caubriere for technical help, and P. Frey for providing material. The authors were supported by grants overseen by the French PIA Lab of Excellence ARBRE (ANR-11-LABX-0002- 01), by the Pôle Scientifique A2F of the Université de Lorraine, by the Région Grand Est (France), by Lorraine Université d’Excellence (LUE) mobility programme DrEAM, by the General Programme of the European Molecular Biology Organization (EMBO Scientific Exchange Grant 9451) and by the INRAE Direction de l’Enseignement Supérieur, des Sites et de l’Europe (DESSE). Research in the Zipfel lab on phytocytokines is supported by funding from the European Research Council under the Grant Agreement No 773153 and by the University of Zurich. Y. Goto was supported by an EMBO Post-Doctoral Fellowship (ALTF 386-2021).

## Author contributions

Conceptualization and design of the study (JL, YG, KB, CZ, BP, SD); data acquisition (JL, GD, EC, RB, BP); data analysis and interpretation (JL, YG, KB, CZ, BP, SD); manuscript drafting (JL, BP). manuscript revision and editing (all authors). All authors contributed to the study and revised, edited, and approved the submitted version.

## Data availability

All the material used in this study is available upon request. All sequences and key information are presented in the Supporting Information Datasets.

