## Supplemental figures for "Genetically-clustered antifungal phytocytokines and receptor proteins function together to trigger plant immune signaling"

### Slide 1
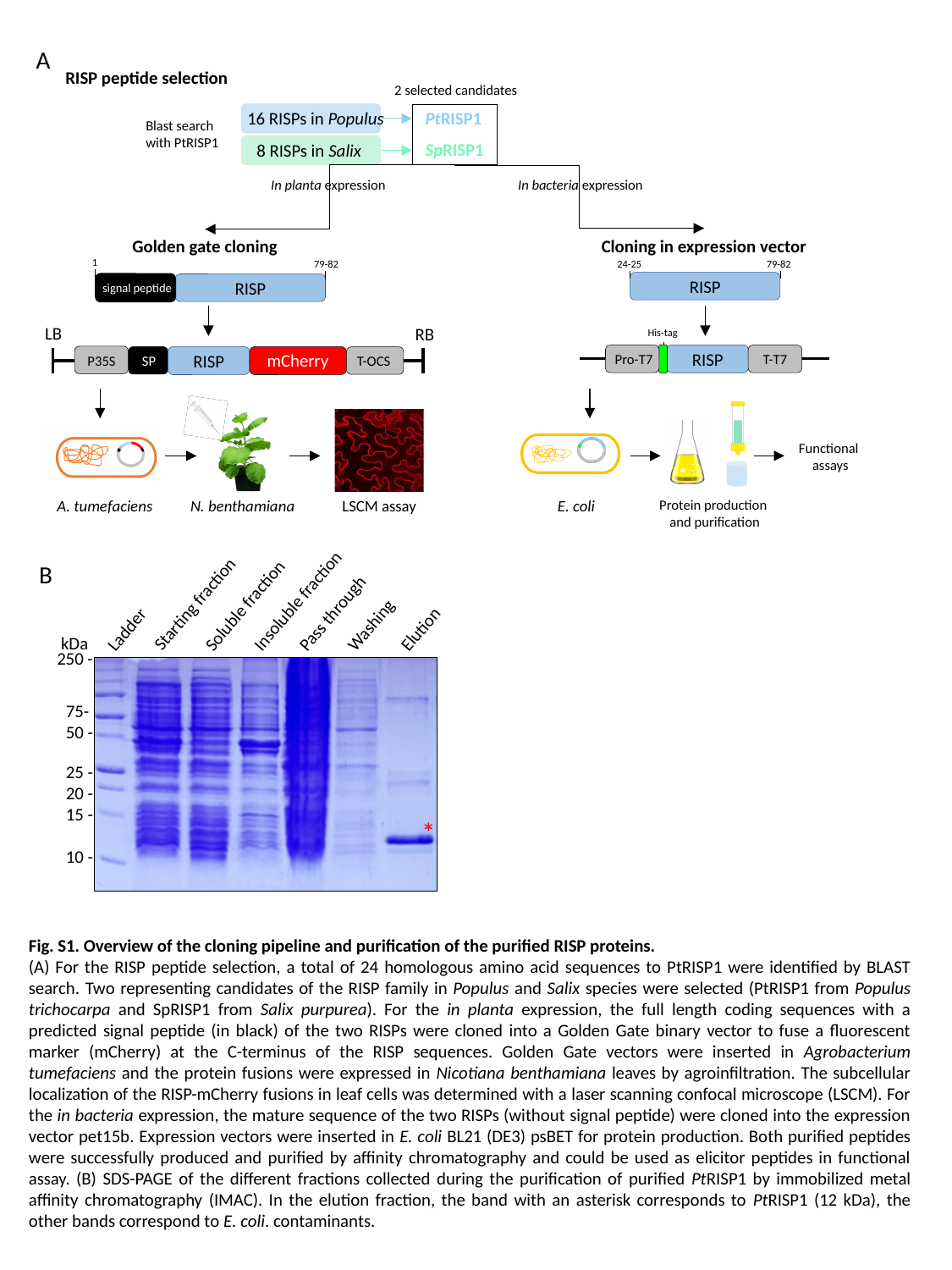

A
RISP peptide selection
2 selected candidates
PtRISP1
SpRISP1
16 RISPs in Populus
Blast search
with PtRISP1
8 RISPs in Salix
In bacteria expression
In planta expression
Golden gate cloning
Cloning in expression vector
1
79-82
signal peptide
RISP
24-25
79-82
RISP
LB
RB
mCherry
P35S
SP
T-OCS
RISP
His-tag
Pro-T7
T-T7
RISP
Functional
 assays
E. coli
Protein production
 and purification
A. tumefaciens
N. benthamiana
LSCM assay
Insoluble fraction
Starting fraction
Soluble fraction
Pass through
Washing
Elution
Ladder
kDa
250 -
75-
50 -
25 -
20 -
15 -
10 -
*
B
Fig. S1. Overview of the cloning pipeline and purification of the purified RISP proteins.
(A) For the RISP peptide selection, a total of 24 homologous amino acid sequences to PtRISP1 were identified by BLAST search. Two representing candidates of the RISP family in Populus and Salix species were selected (PtRISP1 from Populus trichocarpa and SpRISP1 from Salix purpurea). For the in planta expression, the full length coding sequences with a predicted signal peptide (in black) of the two RISPs were cloned into a Golden Gate binary vector to fuse a fluorescent marker (mCherry) at the C-terminus of the RISP sequences. Golden Gate vectors were inserted in Agrobacterium tumefaciens and the protein fusions were expressed in Nicotiana benthamiana leaves by agroinfiltration. The subcellular localization of the RISP-mCherry fusions in leaf cells was determined with a laser scanning confocal microscope (LSCM). For the in bacteria expression, the mature sequence of the two RISPs (without signal peptide) were cloned into the expression vector pet15b. Expression vectors were inserted in E. coli BL21 (DE3) psBET for protein production. Both purified peptides were successfully produced and purified by affinity chromatography and could be used as elicitor peptides in functional assay. (B) SDS-PAGE of the different fractions collected during the purification of purified PtRISP1 by immobilized metal affinity chromatography (IMAC). In the elution fraction, the band with an asterisk corresponds to PtRISP1 (12 kDa), the other bands correspond to E. coli. contaminants.

### Slide 2
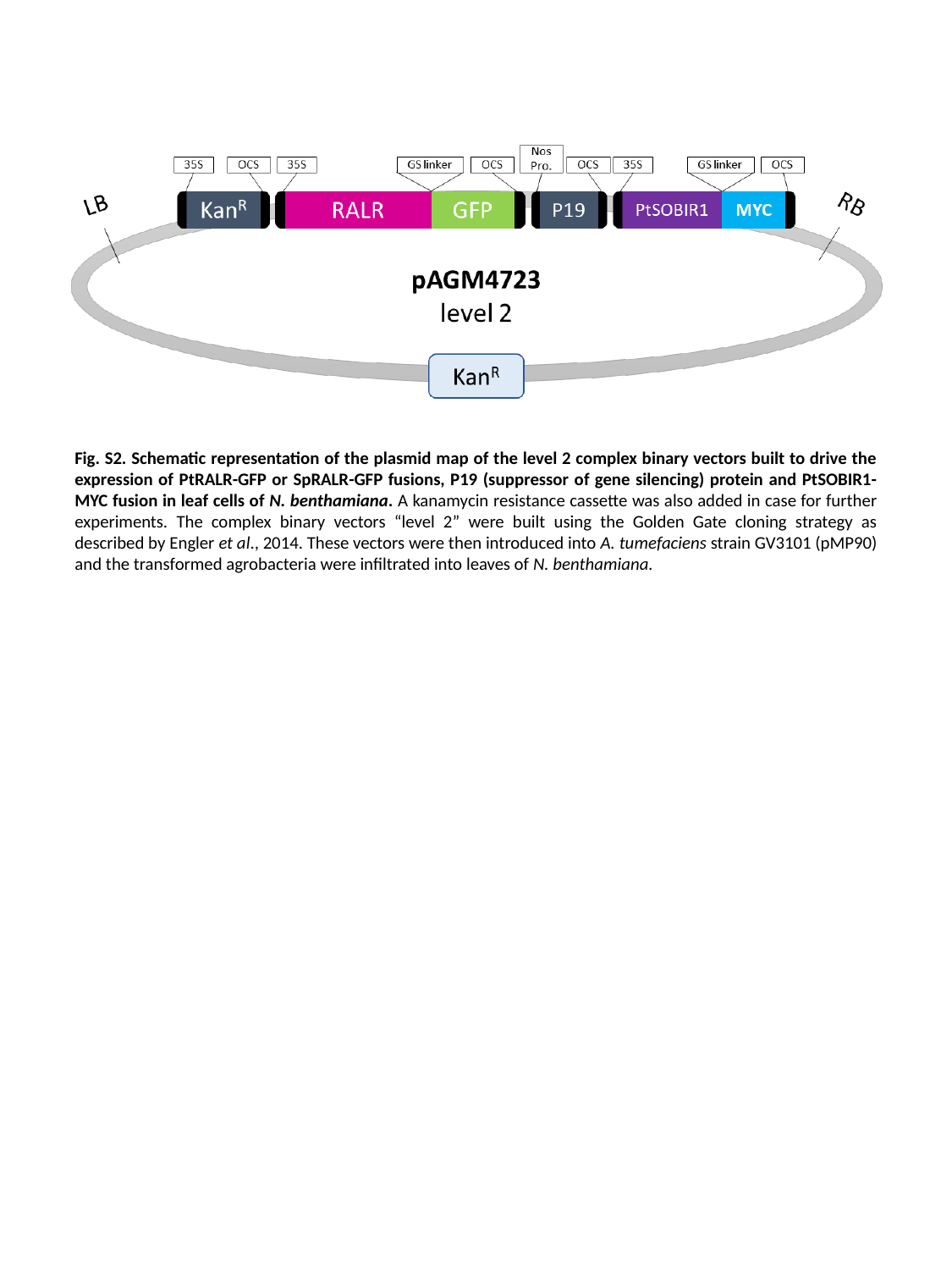

Fig. S2. Schematic representation of the plasmid map of the level 2 complex binary vectors built to drive the expression of PtRALR-GFP or SpRALR-GFP fusions, P19 (suppressor of gene silencing) protein and PtSOBIR1-MYC fusion in leaf cells of N. benthamiana. A kanamycin resistance cassette was also added in case for further experiments. The complex binary vectors “level 2” were built using the Golden Gate cloning strategy as described by Engler et al., 2014. These vectors were then introduced into A. tumefaciens strain GV3101 (pMP90) and the transformed agrobacteria were infiltrated into leaves of N. benthamiana.

### Slide 3
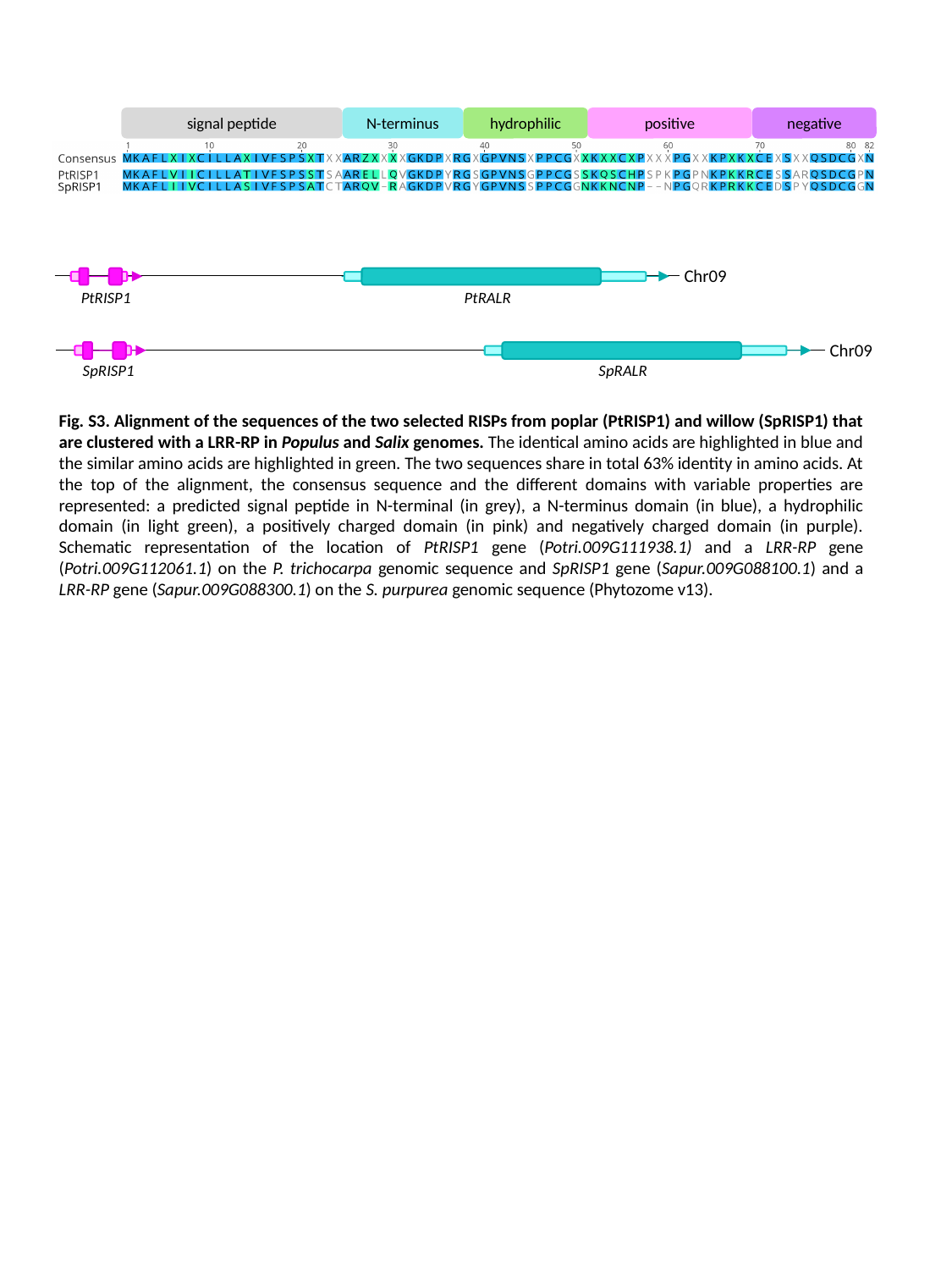

signal peptide
N-terminus
hydrophilic
positive
negative
Chr09
PtRISP1
PtRALR
Chr09
SpRISP1
SpRALR
Fig. S3. Alignment of the sequences of the two selected RISPs from poplar (PtRISP1) and willow (SpRISP1) that are clustered with a LRR-RP in Populus and Salix genomes. The identical amino acids are highlighted in blue and the similar amino acids are highlighted in green. The two sequences share in total 63% identity in amino acids. At the top of the alignment, the consensus sequence and the different domains with variable properties are represented: a predicted signal peptide in N-terminal (in grey), a N-terminus domain (in blue), a hydrophilic domain (in light green), a positively charged domain (in pink) and negatively charged domain (in purple). Schematic representation of the location of PtRISP1 gene (Potri.009G111938.1) and a LRR-RP gene (Potri.009G112061.1) on the P. trichocarpa genomic sequence and SpRISP1 gene (Sapur.009G088100.1) and a LRR-RP gene (Sapur.009G088300.1) on the S. purpurea genomic sequence (Phytozome v13).

### Slide 4
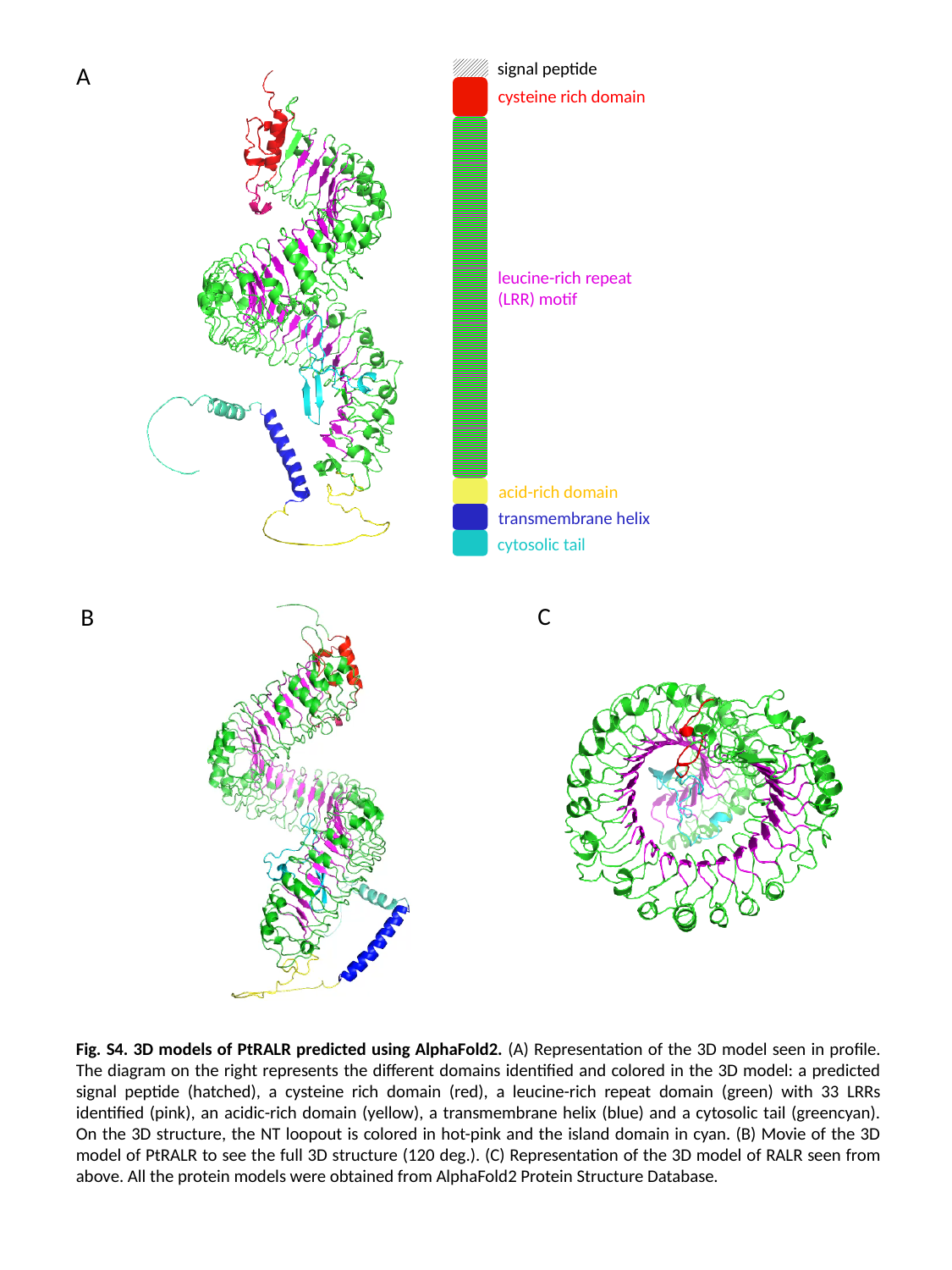

signal peptide
cysteine rich domain
leucine-rich repeat
(LRR) motif
acid-rich domain
transmembrane helix
cytosolic tail
A
C
B
Fig. S4. 3D models of PtRALR predicted using AlphaFold2. (A) Representation of the 3D model seen in profile. The diagram on the right represents the different domains identified and colored in the 3D model: a predicted signal peptide (hatched), a cysteine rich domain (red), a leucine-rich repeat domain (green) with 33 LRRs identified (pink), an acidic-rich domain (yellow), a transmembrane helix (blue) and a cytosolic tail (greencyan). On the 3D structure, the NT loopout is colored in hot-pink and the island domain in cyan. (B) Movie of the 3D model of PtRALR to see the full 3D structure (120 deg.). (C) Representation of the 3D model of RALR seen from above. All the protein models were obtained from AlphaFold2 Protein Structure Database.

### Slide 5
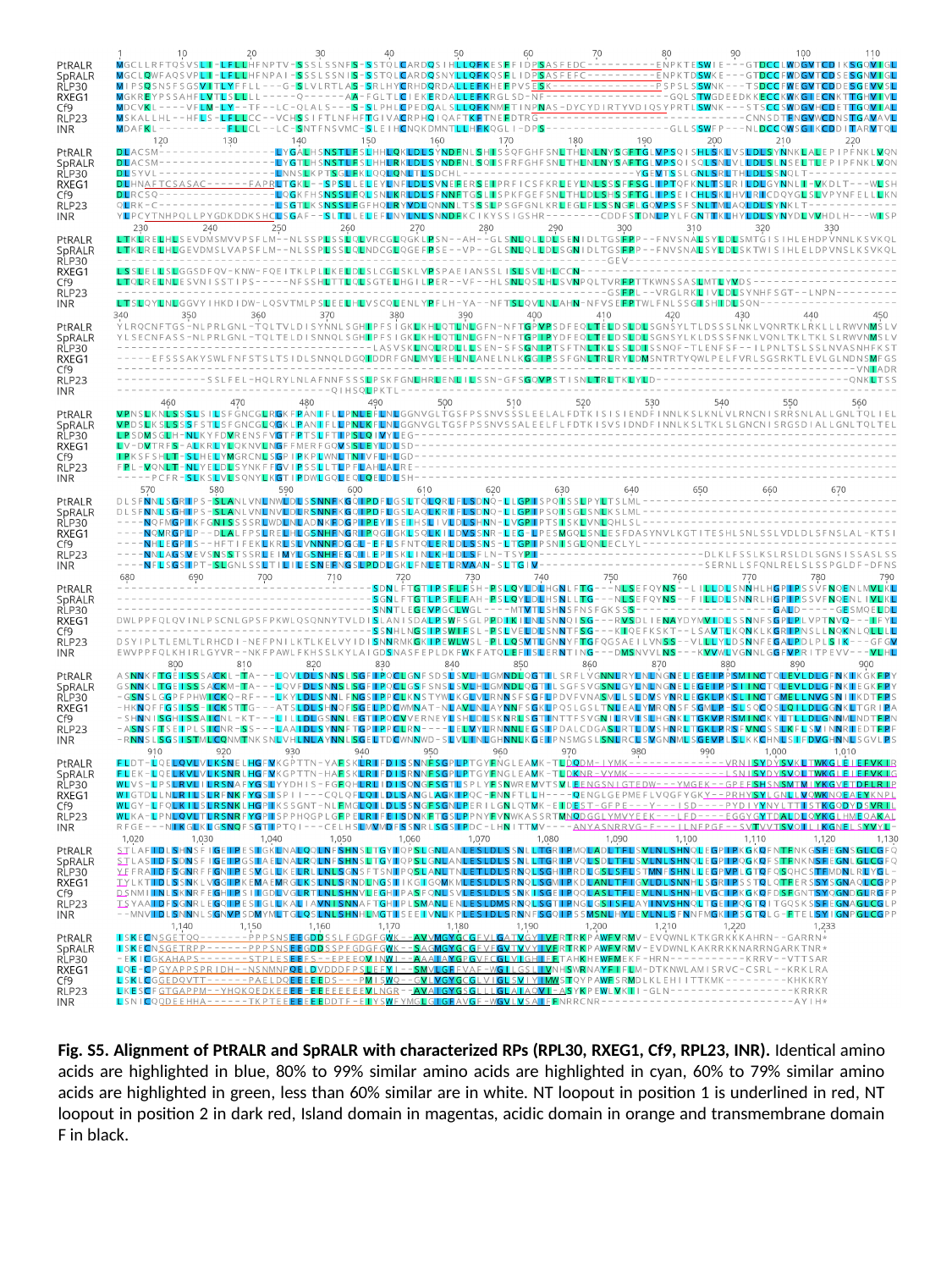

Fig. S5. Alignment of PtRALR and SpRALR with characterized RPs (RPL30, RXEG1, Cf9, RPL23, INR). Identical amino acids are highlighted in blue, 80% to 99% similar amino acids are highlighted in cyan, 60% to 79% similar amino acids are highlighted in green, less than 60% similar are in white. NT loopout in position 1 is underlined in red, NT loopout in position 2 in dark red, Island domain in magentas, acidic domain in orange and transmembrane domain F in black.

### Slide 6
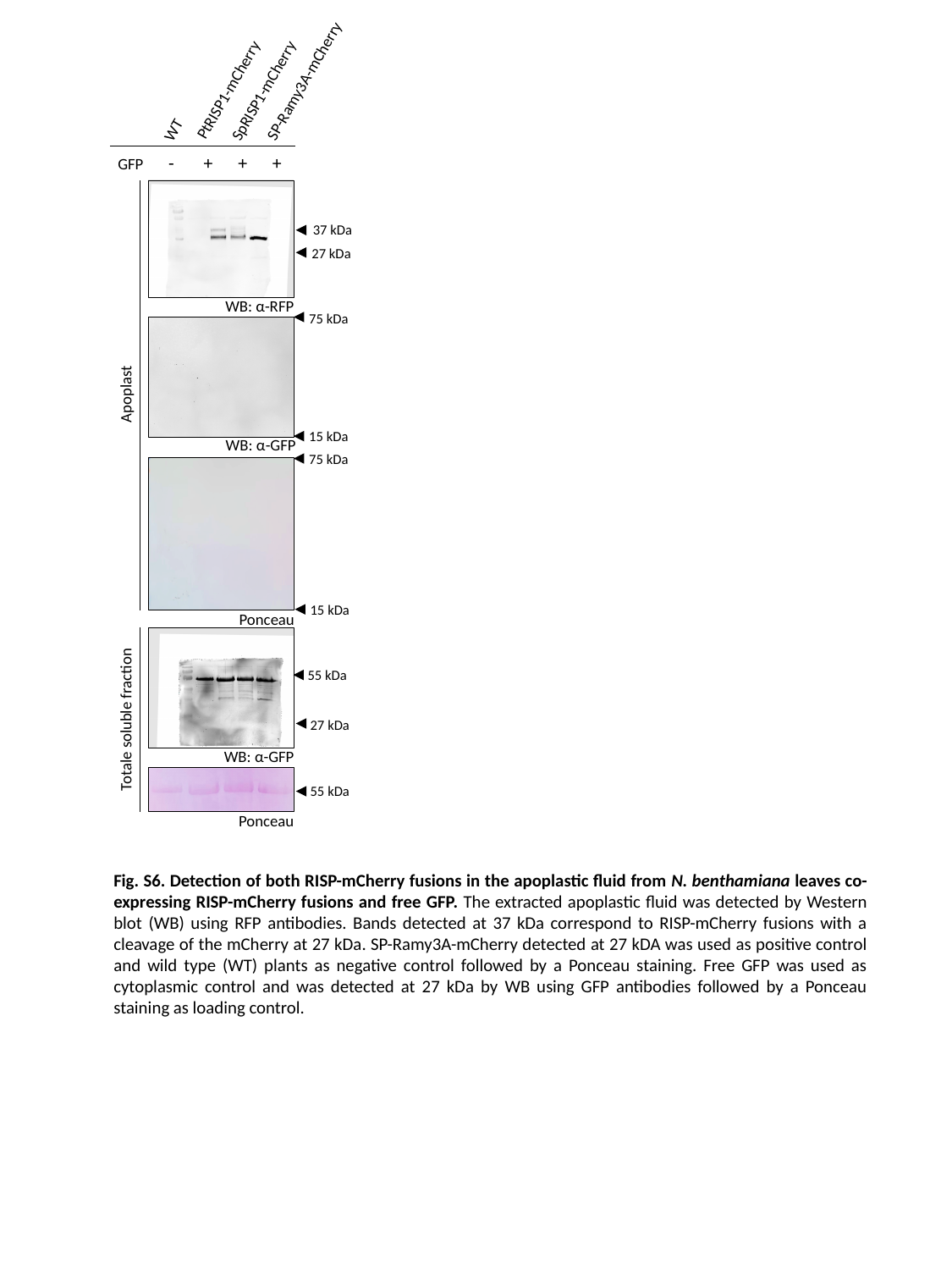

SP-Ramy3A-mCherry
PtRISP1-mCherry
SpRISP1-mCherry
WT
 GFP - + + +
37 kDa
27 kDa
WB: α-RFP
75 kDa
Apoplast
15 kDa
WB: α-GFP
75 kDa
15 kDa
Ponceau
55 kDa
Totale soluble fraction
27 kDa
WB: α-GFP
55 kDa
Ponceau
Fig. S6. Detection of both RISP-mCherry fusions in the apoplastic fluid from N. benthamiana leaves co-expressing RISP-mCherry fusions and free GFP. The extracted apoplastic fluid was detected by Western blot (WB) using RFP antibodies. Bands detected at 37 kDa correspond to RISP-mCherry fusions with a cleavage of the mCherry at 27 kDa. SP-Ramy3A-mCherry detected at 27 kDA was used as positive control and wild type (WT) plants as negative control followed by a Ponceau staining. Free GFP was used as cytoplasmic control and was detected at 27 kDa by WB using GFP antibodies followed by a Ponceau staining as loading control.

### Slide 7
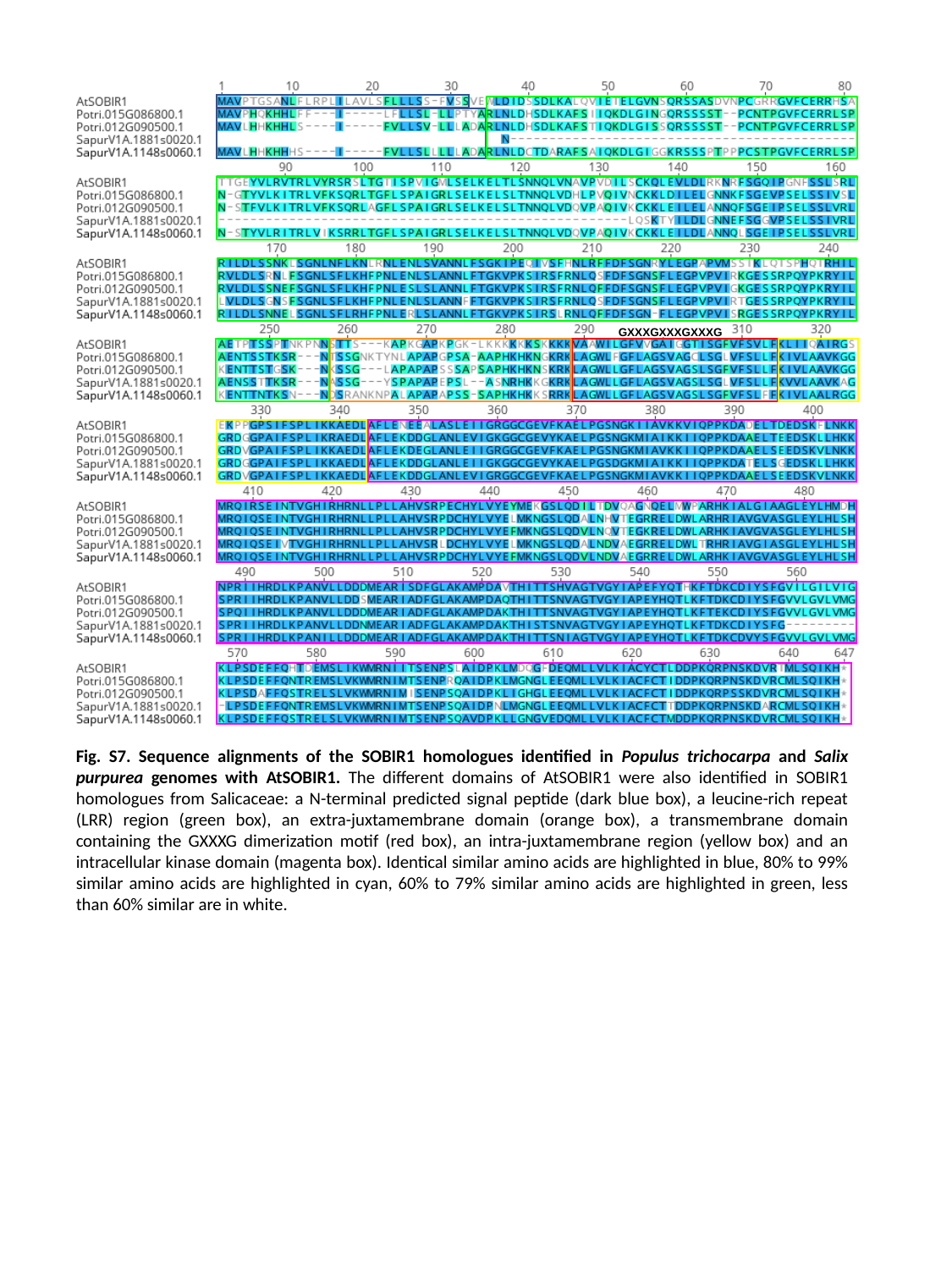

GXXXGXXXGXXXG
Fig. S7. Sequence alignments of the SOBIR1 homologues identified in Populus trichocarpa and Salix purpurea genomes with AtSOBIR1. The different domains of AtSOBIR1 were also identified in SOBIR1 homologues from Salicaceae: a N-terminal predicted signal peptide (dark blue box), a leucine-rich repeat (LRR) region (green box), an extra-juxtamembrane domain (orange box), a transmembrane domain containing the GXXXG dimerization motif (red box), an intra-juxtamembrane region (yellow box) and an intracellular kinase domain (magenta box). Identical similar amino acids are highlighted in blue, 80% to 99% similar amino acids are highlighted in cyan, 60% to 79% similar amino acids are highlighted in green, less than 60% similar are in white.

### Slide 8
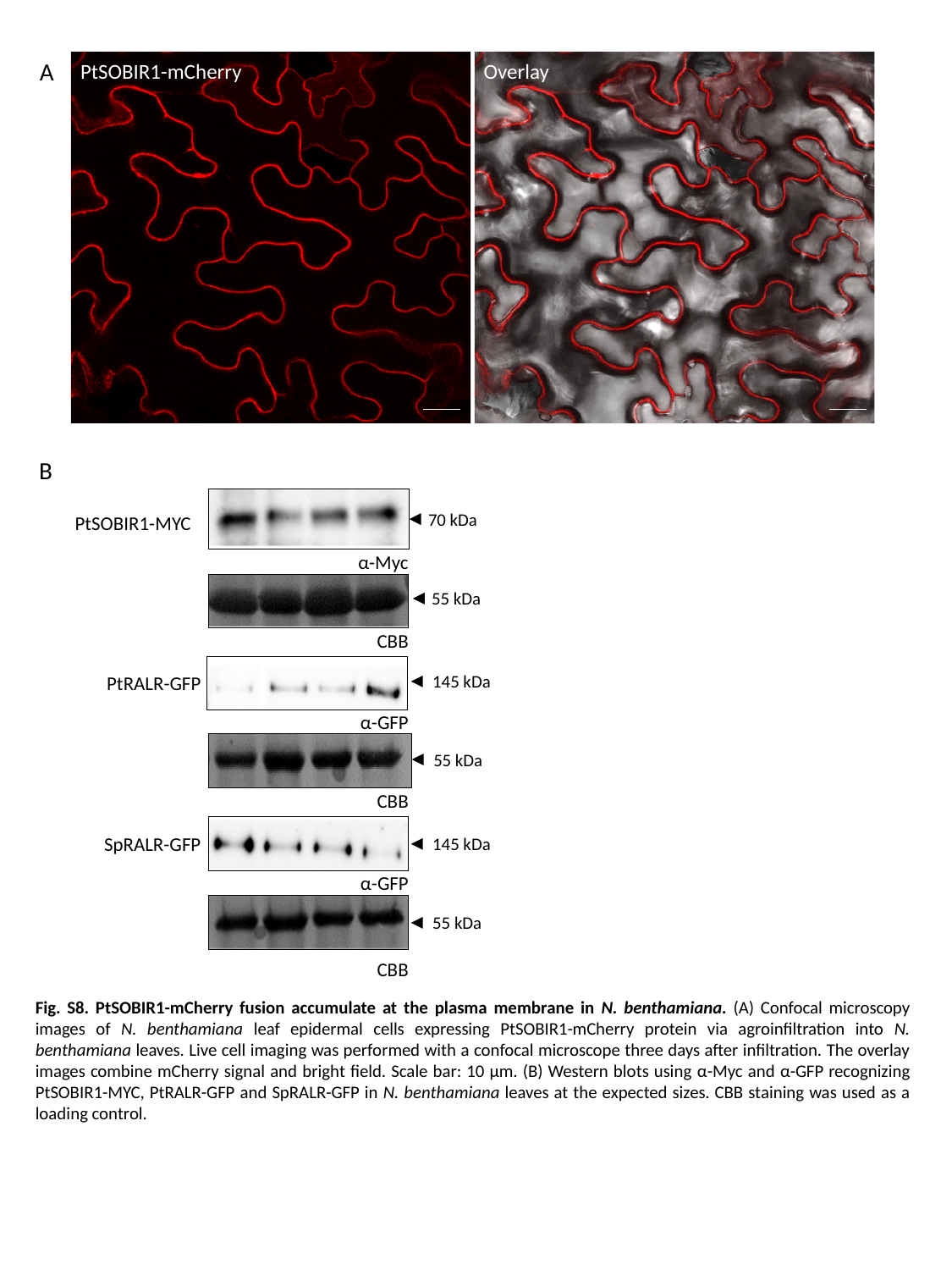

A
PtSOBIR1-mCherry
Overlay
B
70 kDa
PtSOBIR1-MYC
α-Myc
55 kDa
CBB
145 kDa
PtRALR-GFP
α-GFP
55 kDa
CBB
SpRALR-GFP
CBB
145 kDa
α-GFP
55 kDa
CBB
Fig. S8. PtSOBIR1-mCherry fusion accumulate at the plasma membrane in N. benthamiana. (A) Confocal microscopy images of N. benthamiana leaf epidermal cells expressing PtSOBIR1-mCherry protein via agroinfiltration into N. benthamiana leaves. Live cell imaging was performed with a confocal microscope three days after infiltration. The overlay images combine mCherry signal and bright field. Scale bar: 10 µm. (B) Western blots using α-Myc and α-GFP recognizing PtSOBIR1-MYC, PtRALR-GFP and SpRALR-GFP in N. benthamiana leaves at the expected sizes. CBB staining was used as a loading control.
